# THUMPD2 catalyzes *N*^*2*^-methylation on the spliceosome catalytic center of U6 snRNA and regulates pre-mRNA splicing

**DOI:** 10.1101/2023.06.17.545410

**Authors:** Wen-Qing Yang, Jian-Yang Ge, Xiao-Feng Zhang, Lin Lin, Wen-Yu Zhu, Yi-Gong Shi, Bei-Si Xu, Ru-Juan Liu

## Abstract

How the relatively evolutionarily conserved spliceosome is able to manage the enormously expanded number of splicing events that occur in humans (∼200,000 vs. ∼400 reported for yeast) is not well understood. Here, we show deposition of one RNA modification-N2-methylguanosine (m^2^G)-on the G72 nucleoside of U6 snRNA (known to function as the catalytic center of the spliceosome) results in profoundly increased pre-mRNA splicing activity in human cells. This U6 m^2^G72 modification is conserved among vertebrates. Further, we demonstrate that THUMPD2 is the methyltransferase responsible for U6 m^2^G72 and show that it interacts with an auxiliary protein (TRMT112) to specifically recognize both sequence and structural elements of U6. *THUMPD2* KO blocks U6 m^2^G72 and down-regulates the pre-mRNA splicing activity of major spliceosome, yielding thousands of changed alternative splicing events of endogenous pre-mRNAs. Notably, the aberrantly spliced pre-mRNA population of the *THUMPD2* KO cells elicits the nonsense-mediated mRNA decay (NMD) pathway and restricts cell proliferation. Our study thus demonstrates how an RNA epigenetic modification of the major spliceosome differentially regulates global pre-mRNA splicing.

## Main text

Splicing of precursor messenger RNA (pre-mRNA) plays a pivotal role in eukaryotes, which is performed by a mega RNA-protein complex called spliceosome (*1*). From yeast to mammals, the frequency of splicing events increases from 8% to around 98% of all protein-coding genes. There are approximately 400 genes containing a single intron in about 5,000 total genes in *Saccharomyces cerevisiae*, while there are approximately 20,000 genes, which on average contain 8 introns in humans (*2*). Intriguingly, plenty of biochemical and structural studies show that the overall organization of spliceosome and the configuration of the splicing catalytic core are extremely conserved between human and yeast (*2, 3*). The enormously expanded splicing events in higher eukaryotes are therefore a challenge to the operational capability of spliceosome, but the precise underlying mechanism remain unclear. The spliceosome is a protein-orchestrated ribozyme dynamically formed anew on pre-mRNA by five conserved small nuclear RNAs (snRNAs: U1, U2, U4, U5, and U6) with multitudinous proteins (*4*). The splicing process is proven to be catalyzed by snRNAs, and U6 locates in the splicing catalytic core of spliceosome (*5*). (Fig. S1a). The Internal Stem Loop (ISL) of U6 constitutes a major part of splicing catalytic center (*6*). During pre-mRNA splicing, nucleotides from the splicing catalytic center chelate two catalytic metal ions to fulfil branching and exon ligation (*7*).

RNA modifications play key regulatory role in various biological events (*8*). U6 is heavily modified post-transcriptionally in higher eukaryotes (*9-11*), with pseudouridylation (ψ) and 2’-*O*-methylation (Nm) in multiple sites and two individual *N*^*6*^-methyladenosine (m^6^A, nucleoside 43 in human U6) and *N*^*2*^-methylguanosine (m^2^G, nucleoside 72 in human U6) (*11-13*). The ψ and Nm of U6 were predominantly introduced by Box C/D snoRNP (dyskerin) and Box H/HCA snoRNP (fibrillarin), respectively (*5*) (Fig. 1a and Fig. S1b), and were proposed to increase base-stacking and enhance base-pairing in U6 (*11*). Recently, U6 m^6^A was found to be vital for recognizing 5’ splice site, and METTL16 was identified as its methyltransferase (*14, 15*). However, the role of the only other base methylation-U6 m^2^G72 remains elusive. Structures of human major spliceosome reveal that G72 in the ISL is involved in forming the activated catalytic center of spliceosome and directly binds catalytic ion M1 (*7*) (Fig. S1c). In the two-metal-ion mechanism model for splicing catalysis (*16, 17*), human U6-G72, -U74, and pre-mRNA provide Sp phosphate oxygen to bind two metal ions which bond with guanosine’s phosphate of 5’ splice site to excise intron (*7*). In some sequenced U6 from mammals, an m^2^G modification exhibits on G72. Although this modification was firstly identified four decades ago (*12, 13*), its impacts on pre-mRNA splicing by spliceosome as well as its physiological roles remain unclear, largely because the responsible modifying enzyme has not been identified.

**Fig. 1.**
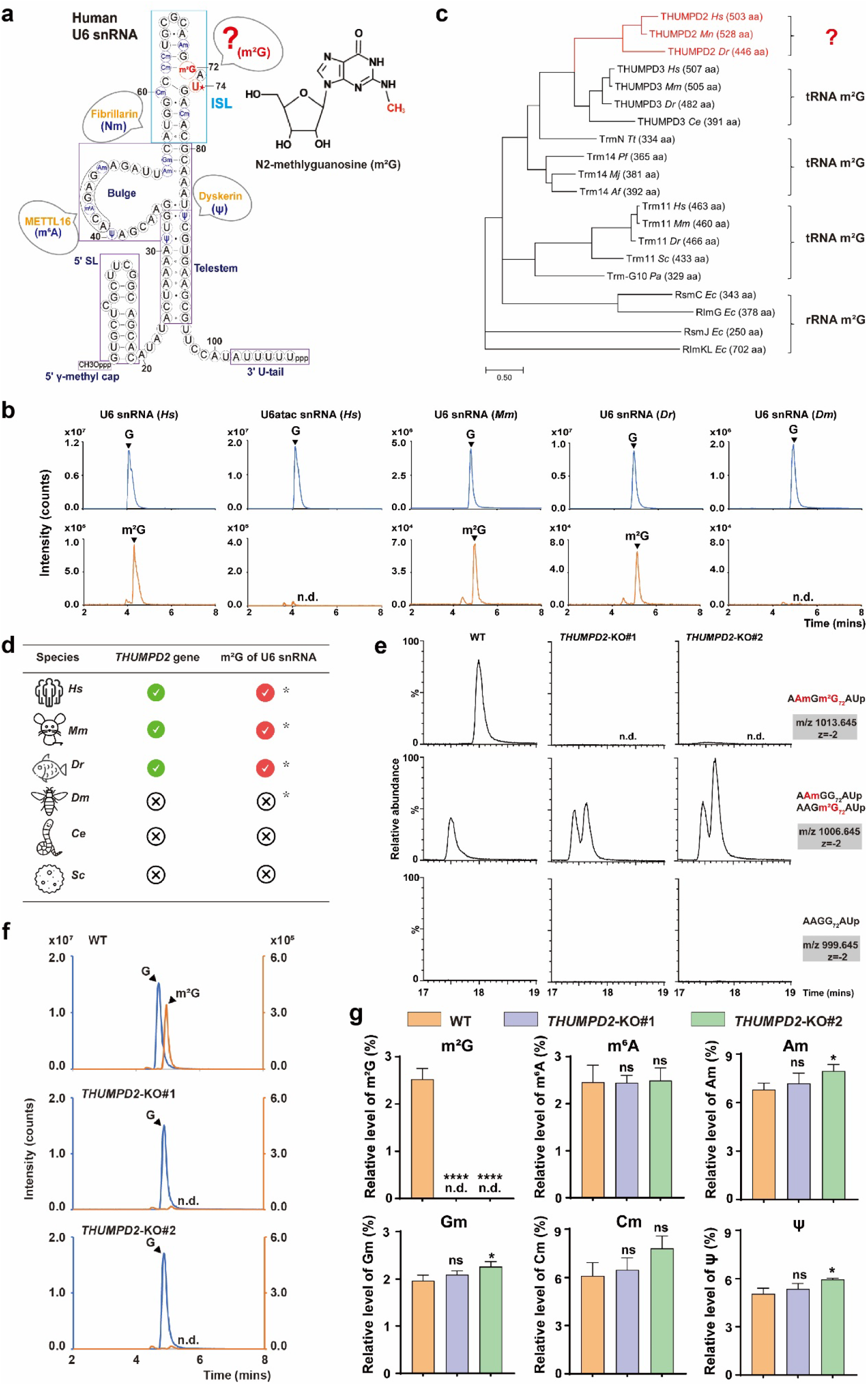
Analysis of U6 m^2^G72 in eukaryotic model organisms and identification of THUMPD2 as the responsible methyltransferase. (a) Secondary structures of U6 snRNA from *Homo sapiens* marked with the following elements: 5’γ-methyl cap, 5’SL, ISL, Bulge, Telestem, and 3’U-tail. All the identified RNA modifications are labeled, and responsible modifying enzymes are listed, respectively. The chemical structure of m^2^G is labelled on the upper right. (b) Mass chromatograms of nucleosides, G (Q1/Q3 = 284.1/152.2) (upper panel) and m^2^G (Q1/Q3 = 298.1/166.1) (lower panel) of U6 extracted from human HEK293T cells, brain from 11-weeks mouse, tail from zebrafish, and whole drosophila, respectively. And U6atac snRNA extracted from HEK293T cells was also analyzed. The abbreviations used: *Hs, Homo sapiens*; *Mm, Mus musculus*; *Dr, Danio rerio*; and *Dm, Drosophila melanogaster*. (c) Phylogenetic analysis of RNA m^2^G enzymes indicates that THUMPD2 is likely an evolutionarily conserved N2-methylation-related enzyme in vertebrates. aa, amino acids. (d) The evolution of *THUMPD2* genes and m^2^G at U6 from yeast to human. * Means m^2^G of U6 is verified in this study. The abbreviations used: *Ce, Caenorhabditis elegans* and *Sc, Saccharomyces cerevisiae*. (e) RNA-fragment-MS analysis of m^2^G-containing fragments of U6 snRNA, extracted from WT and both two *THUMPD2* KO HEK293T cells, digested by RNase A with (upper two panels) or without (lower panel) m^2^G. The sequences of modified and unmodified fragments are indicated. The m/z value and charge state of each fragment are shown on the right. m^2^G, N2-methylguanosine. (f) Mass chromatograms of nucleosides, G and m^2^G of U6 extracted from WT and both two *THUMPD2* KO HEK293T cells. Target peaks are indicated by black asterisk. The relative abundance of G is used as control. G, guanosine. (g) Quantification of m^2^G, m^6^A, Am, Gm, Cm, and ψ level of U6 between WT and both two *THUMPD2* KO HEK293T cells. Data were normalized to relative abundance of A. Data information: In g, statistical analysis was performed using t-tests, and the error bars indicate mean ± SD for three independent experiments. n.d., not detected; ns, not statistically significant; ∗*p* < 0.05; ∗∗∗∗*p* < 0.0001.

Here we show that the m^2^G72 is present in U6 of zebrafish, mouse, and human, but is absent in U6atac-the functional analog of U6 in minor spliceosome. Using a reverse genetics approach coupled with RNA-mass spectrometry, we reveal that THUMP domain containing protein 2 (THUMPD2) as its methyltransferase. THUMPD2 interacts with an auxiliary protein, TRMT112, to catalyze U6 m^2^G72 formation. *In vitro* pre-mRNA splicing assays showed that loss of U6 m^2^G72 down regulates the activity of major spliceosome. In *THUMPD2* KO human cells, loss of U6 m^2^G72 causes thousands of changed alternative splicing events and up regulates the expression of genes in nonsense-mediated mRNA decay (NMD) pathway.

### Analysis of U6 m^2^G72 and identification of THUMPD2 as the responsible methyltransferase

A m^2^G modification was identified on the splicing active site (G72) of U6 snRNA in some mammals, G72 directly participates in pre-mRNA processing via binding catalytic ion M1 (*16, 17*). However, the function of U6 m^2^G72 remain unclear as well as the responsible writer. To verify U6 m^2^G72 in some representative eukaryotes, we purified U6 snRNAs from human, mouse, zebrafish, and drosophila, and profiled RNA modifications using RNA mass spectrometry (RNA-MS) (Fig. S3a). A m^2^G modification of U6 was evident for the three examined vertebrates (human, mouse, and zebrafish) but not for drosophila (Fig. 1, b and d, and Fig. S3c). The m^2^G was indeed at position 72 of U6 isolated from human HEK293T cells was further confirmed by the mass spectrometry of RNA fragments (RNA-fragment-MS). The fragments containing m^2^G72 (AAmGm^2^G_72_AU and AAGm^2^G_72_AU) were detected as the majority (70%-100%) comparing to G72 unmodified fragments (AAmGG_72_AU and AAGG_72_AU) (Fig. 1e left panel), suggesting a high level of m^2^G72 modification of U6 *in vivo*. U6atac is the U6 functional analog in the minor spliceosome, and folds into a similar secondary structure as U6 (*18*); the equivalent nucleoside G72 (G44 in human U6atac) also presents in the splicing catalytic center (Fig. S3b). We found that U6atac is not modified with m^2^G either in human, mouse, or zebrafish (we could not obtain U6atac from zebrafish although we made lots of efforts to purify it from several zebrafish, which might attribute to infinitesimal U6atac in zebrafish) (Fig. 1b and Fig. S3d). These findings suggested that m^2^G is a prevalent modification on U6 snRNA in the examined vertebrates.

To identify the writer for the U6 m^2^G72, we performed a reverse genetics approach coupled with RNA-MS. First, we used a phylogenetic analysis of known and predicted RNA m^2^G methyltransferases and revealed three candidates in vertebrates (Fig. 1, c and d). Among them, we focused on THUMPD2-a predicted RNA m^2^G methyltransferase, because it is the only one conserved in vertebrates but not in drosophila or lower eukaryotes revealed by protein sequence alignments (Fig. S2). We first confirmed that THUMPD2 was predominantly located in the nucleus where U6 snRNA stays by Immunofluorescence assays (Fig. S6a). We therefore knocked out *THUMPD2* in human HEK293T, HeLa and HeLa S3 cells using the CRISPR-Cas9 system (Fig. S4, a, b and c). The U6 snRNA was then isolated from *THUMPD2* KO cells and subjected to RNA-MS and RNA-fragment-MS. Notably, m^2^G72 in U6 was completely lost in all the *THUMPD2* KO HEK293T cells (Fig. 1e and 1f), but the level of the rest modifications including m^6^A, Ψ, and Nm (Am, Gm, and Cm) remained unchanged (Fig. 1g and Fig S5). When *THUMPD2* KO HEK293T cells were obviously rescued by plasmid encoded THUMPD2, and m^2^G72 was efficiently restored, demonstrating that THUMPD2 is responsible for the methylation of U6 m^2^G72 (Fig. S3e middle panel).

The function of THUMPD2 has barely been characterized before, we thus measured its expression pattern in human cell lines and mouse tissues. It is widely expressed in all the tested human cell lines derived from embryonic kidney, neuroblastoma, liver, cervix uterus, tibia, breast, lung, and retina, respectively (Fig. S7a). *Thumpd2* mRNA is expressed in all the tested tissues with a relative high level in thymus, brain, and eye from 11-week-old mice (Fig. S7b).

In summary, U6 m^2^G72 is conserved in vertebrate and its writer THUMPD2 is widely expressed in various cells and tissues.

### Molecular basis of THUMPD2-mediated U6 m^2^G72 modification

Through our efforts to use *in vitro* methylation assays with purified THUMPD2 protein to validate its methyltransferase activity, we noted that standalone THUMPD2 showed no detectable activity (Fig. 2, a and b). Since many RNA methyltransferases function as a complex for their catalysis, this prompted us to identify THUMPD2-binding proteins (*19-21*). We thus performed immunoprecipitation coupled with proteomic analysis using THUMPD2-HA and found TRMT112 (Fig. S6b and Table S1). TRMT112 is conserved in higher eukaryotes and known to bind and regulate other methyltransferases in methylation of rRNAs, tRNAs, and proteins (*22, 23*) (Fig. S8). The interaction between THUMPD2 and TRMT112 in HEK293T cells was confirmed by immunoprecipitation assays (Fig. S6, c, d, and e). The direct interaction between THUMPD2 and TRMT112 from human, mouse, and zebrafish was confirmed by gel filtration analysis using purified proteins in a high-salt concentration buffer (500 mM NaCl). The chromatography results showed that THUMPD2 and TRMT112 formed a stable binary complex (Fig. S9a). We further generated a structural model of human THUMPD2-TRMT112 (Fig 2, a and c), in which THUMPD2 contains a classical Rossman-Fold methyltransferases domain linked to a THUMP containing domain, while TRMT112 has a small single domain. *In vitro* methylation assays showed that THUMPD2-TRMT112 complex of all the three tested species could catalyze m^2^G methylation to their cognate U6 snRNA (*24*) (Fig. 2d and Fig. S9, b and c). The methylation site G72 was confirmed by RNA-fragment-MS and mutagenesis studies *in vitro* (Fig. S6, f and g). These results indicate that THUMPD2 forms a complex with TRMT112 to catalyze U6 m^2^G72 formation.

**Fig. 2.**
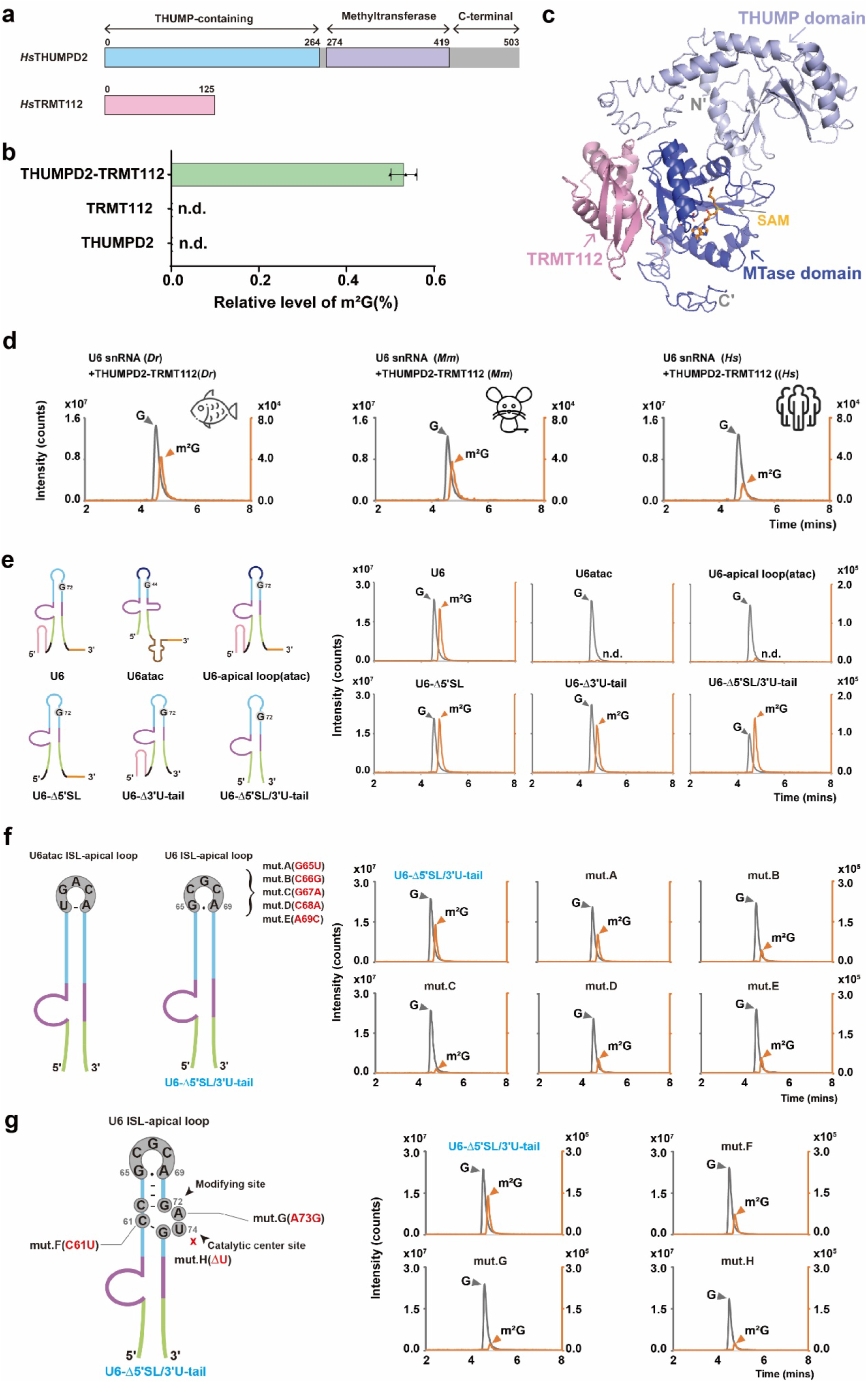
THUMPD2 interacts with TRMT112 to catalyze m^2^G72 by recognizing U6 specific sequence and structural motifs. (a) Domain composition of THUMPD2 and TRMT112 of human. *Hs*THUMPD2 contains 503 aa and *Hs*TRMT112 contains 125 aa. THUMPD2 contains two main domains: an ancient RNA-binding THUMP domain followed by a S-adenosylmethionine (SAM)-dependent MTase domain. (b) Quantification of the methylation activity of purified *Hs*THUMPD2, *Hs*TRMT112, and *Hs*THUMPD2-TRMT112 complex to U6. Data were normalized to the relative abundance of G. (c) The structural model of the *Hs*THUMPD2-TRMT112 complex was generated by a protein structure homology-modelling server combined with manual structural superimposition and docking. The model is shown in cartoon form, and the MTase (dark blue) and THUMP domain (lilac) of THUMPD2 with SAM (orange) plus TRMT112 (pink) are indicated in different colors. (d) Mass chromatograms of G and m^2^G from U6 of corresponding species after incubation with the purified THUMPD2-TRMT112 complex from zebrafish, mouse, and human, respectively. Target peaks are indicated by grey and orange triangle, respectively. (e) Elements on U6 that are recognized by THUMPD2-TRMT112. Mass chromatograms of G and m^2^G from U6, U6atac, and U6 mutants after incubation with THUMPD2-TRMT112. Diagram of secondary structures of different truncated or substitutive forms of U6 were showed on the left. (f) Mutations on the ISL-apical loop. Diagram of different primary sequence of ISL-apical loop between U6atac and U6 are showed on the left. The distinguishing nucleoside of U6 was mutated into the corresponding base of U6atac of ISL-apical loop. And the identical nucleoside of U6 that C68 and A69 were mutated into A68 or C69, respectively. From 5’ to 3’ the mutations were successively named as m.A, m.B, m.C, m.D, and m.E. Mass chromatograms of G and m^2^G of U6 telestem and different mutants after incubation with THUMPD2-TRMT112. (g) Diagram of detailed primary sequence around modifying site G72 of U6 were showed on the left. m.F, in which C61 was mutated into U61; m.G, which A73 was mutated into G73; m.H, in which the catalytic site U74 was deleted. Mass chromatograms of G and m^2^G of different U6 mutants after incubation with THUMPD2-TRMT112. In Fig. d, e, f, and g, target peaks are indicated by grey (G) and orange (m^2^G) triangle. Data information: In b, statistical analysis was performed using t-tests, and the error bars indicate mean ± SD for three independent experiments. n.d., not detected.

THUMPD2-TRMT112 catalyze m^2^G formation on U6, but not U6atac (Fig. 1b and Fig. S3, c and d). To elucidate the molecular mechanism of substrate specificity of THUMPD2-TRMT112, we made mutagenic analysis based on differences between U6 and U6atac followed by *in vitro* methylation assays. U6 and U6atac are different in the 5’ Stem Loop (5’SL), 3’ U-tail, and apical loop of ISL (5) (Fig.1a and Fig. S3b). We truncated 5’SL and/or 3’ U-tail or substituted the ISL-apical loop on U6 by the U6atac counterpart to generate four U6 variants, and the *in vitro* methylation assays by THUMPD2-TRMT112 showed that m^2^G formation of U6 were unaffected by truncation of 5’SL or 3’ U-tail but was completely lost when substituted the ISL-apical loop to that of U6atac (Fig. 2e). Subsequently, we mutated each nucleoside on U6 ISL-apical loop (_65_GCGCA_68_), and the *in vitro* methylation assays showed that the m^2^G level of all the mutants methylated by THUMPD2-TRMT112 decreased to ∼15-50% in comparison to control sample (Fig. 2f). These results indicate that the _65_”GCGCA”_68_ motif of U6 ISL-apical loop is required for recognition by THUMPD2-TRMT112.

G72 involves in the formation of RNA splicing catalytic center. The nucleosides (U74 and A73) in the vicinity of G72 forms a specific bulged structure (*25, 26*), in which G72 base pairs with C62 (Fig. 1a and Fig. S10). Since the methyl group of m^2^G is on the purine base, we assumed that the purine base of G72 must flip out to be modified by THUMPD2. Pursuing the involvement of this bulged structure during m^2^G72 deposition, we mutated C61 to U which will form a Watson-Crick base pair with A73, or mutated A73 to G that might form G73-C61 base pair, or directly deleted U74 to disrupt the bulged structure. The *in vitro* methylation assays showed that the m^2^G level of all these mutants methylated by THUMPD2-TRMT112 decreased to ∼7-25% in comparison to control sample (Fig. 2g), indicating that the specific bulged structure around G72 is essential for recognition of U6 by THUMPD2-TRMT112.

In summary, THUMPD2 interacts with TRMT112 to catalyze U6 m^2^G72 formation by recognizing the _65_”GCGCA”_68_ motif of the U6 ISL-apical loop and the specific bulged structure near G72.

### Loss of U6 m^2^G72 decreases the pre-mRNA splicing activity of spliceosome *in vitro*

In human major spliceosome, U6 G72 involves in forming the activated catalytic center and functions in pre-mRNA splicing by chelating a catalytic metal ion M1 that firstly attacks 5’ splice site for cleavage and next binds to 3’ splice site for exon ligation (*16*) (Fig. S1c). Since G72 participates in pre-mRNA splicing process, we sought to investigate whether m^2^G modification on G72 has any impact in the pre-mRNA splicing activity of spliceosome using *in vitro* splicing assays.

We first confirmed that *THUMPD2* KO does not alter the U6 level in all the tested HEK293T, HeLa, and HeLa S3 cells (Fig. 3a). We subsequently leverage *in vitro* pre-mRNA splicing assays using different HeLa S3 nuclear extracts. An intron-containing MINX RNA (with two exons interrupted by an intron with canonical splice sites of the major spliceosome with “GU-AG” rule) was incubated with HeLa S3 nuclear extracts. Un-spliced and spliced MINX RNA was analyzed with RT-PCR at varying time points from 10-90 minutes to assess pre-mRNA splicing (Fig. 3b). The amount of spliced MINX RNA in both two *THUMPD2* KO cells was obviously reduced compared to WT cells starting at the 20 minutes time point, and the extent of the difference became apparently larger with increased reaction time (Fig. 3c). Notably, the pre-mRNA splicing activity could be rescued by nuclear extract supplemented with wild type THUMPD2 obviously, but not with enzymatically inactive THUMPD2 mutant D329A (Fig. 3d and S3e right panel). These results indicate that splicing activity of MINX RNA is obviously impaired upon *THUMPD2* KO, and the regulation of splicing activity is dependent on the U6 m^2^G72 methylation capability of THUMPD2.

**Fig. 3.**
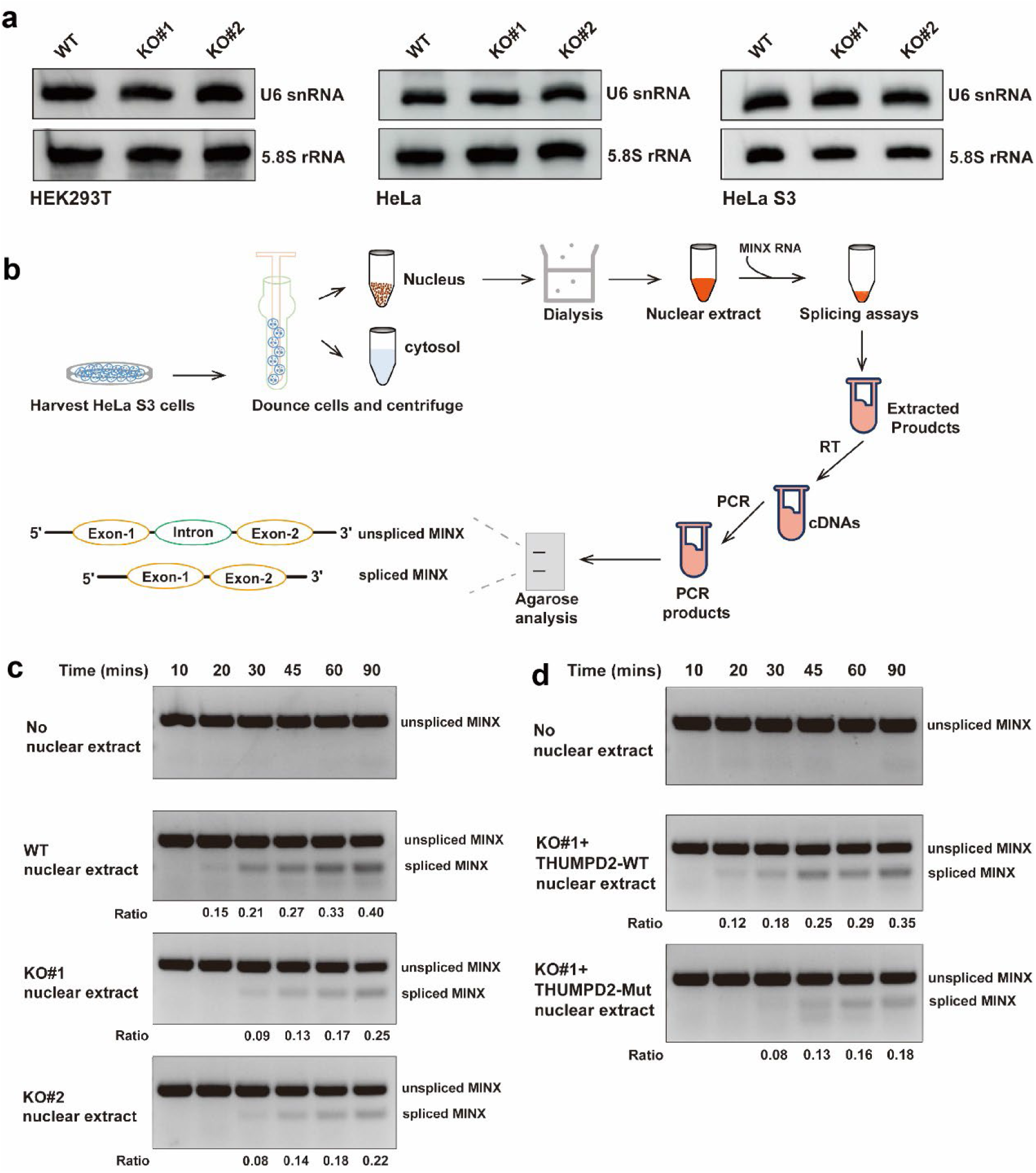
Analysis *in vitro* splicing activity assays in *THUMPD2* KO cells. (a) Northern blotting analysis of the steady state level of U6 snRNA (upper lane) between WT and both two *THUMPD2* KO HEK293T, HeLa, and HeLa S3 cells. 5.8S rRNA (lower lane) were used as control. (b) Schematic of the *in vitro* splicing assays using HeLa S3 nuclear extract. (c) Detection of splicing velocity of MINX RNA in WT and both two *THUMPD2* KO HeLa S3 cells by RT-PCR. The un-spliced (upper lane) and spliced (lower lane) MINX RNA are detected in varying time points from 10 to 90 minutes. (d) Detection of splicing velocity of MINX RNA in *THUMPD2* KO#1 HeLa S3 cells rescued by plasmid-encoded human wild type THUMPD2 and enzymatically inactive THUMPD2 (D329A) by RT-PCR. The un-spliced (upper lane) and spliced (lower lane) MINX RNA are detected in varying time points from 10 to 90 minutes. The Ratio in c and d represents un-spliced MINX in the upper lane and spliced MINX in the lower lane. visualization of RT-PCR products using Gelred-stained agarose gels.

Thus, our data show that the pre-mRNA splicing activity of spliceosome is downregulated when loss of U6 m^2^G72 modification by *THUMPD2* KO.

### *THUMPD2* KO causes altered splicing of endogenous pre-mRNAs

We next ask whether the *THUMPD2* KO could impact endogenous pre-mRNA substrates across the transcriptome. To this end, we performed RNA-seq analysis of WT and both two *THUMPD2* KO HEK293T cells to assess global pre-mRNA splicing changes and used a junction read and exon expression approach rMATS (*27*). We first called All KO samples (4 replicates) versus WT (2 replicates). To assure the reproducibility of results, we also called KO1 or KO2 (each with two replicates) versus WT independently and filter the results by excluding those were not significant (*p* < 0.05) in either KO1 or KO2. At last, we excluded low count events (> 10 junction reads, > 0.5 counts per million) to increase the confidence. A total of 4113 significant alternative splicing (AS) events were evident in the *THUMPD2* KO cells, including 1907 skipped exons (SE), 906 alternative 3’ splice sites (A3SS), 604 retained introns (RI), 407 mutually exclusive exons (MXE), and 289 alternative 5’ splice sites (A5SS) (Fig. 4a), affecting a total of 1923 genes (|ψ|≥5%, false discovery rate [FDR]≤0.05, > 10 junction reads, > 0.5 counts per million).

**Fig. 4.**
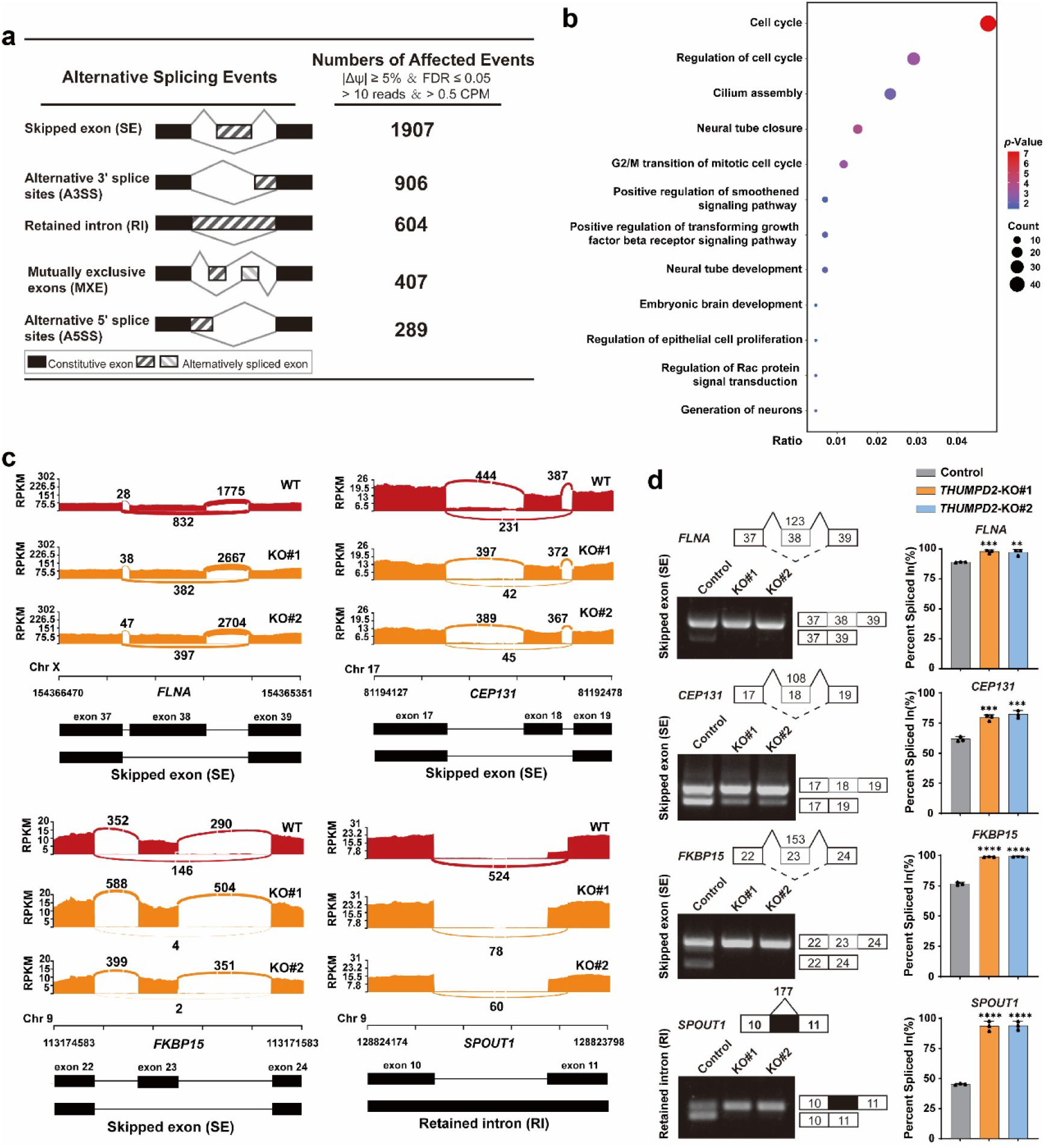
Analysis of endogenous pre-mRNA splicing in *THUMPD2* KO cells. (a) Summary of the different alternative splicing (AS) events identified in *THUMPD2* KO HEK293T cells and the numbers of splicing events affected by *THUMPD2* depletion. (b) Bubble chart that shows the up or down regulated skipped exon (SE) events of protein-coding genes in *THUMPD2* KO HEK293T cells showing enriched GO terms in the biological process part. (c) Sashimi plots of RNA-seq junction reads counts for representative skipped exon (SE) events in *FLNA, CEP131*, and *FKBP15* and retained intron (RI) event in *SPOUT1* in both two *THUMPD2* KO HEK293T cells. RPKM, reads per kilobase per million mapped reads. (d) Validation of SE events in *FLNA, CEP131*, and *FKBP15* and RI event in *SPOUT1* in both two *THUMPD2* KO HEK293T cells by RT-PCR with β-actin serving as a loading control. Left: visualization of RT-PCR products using Gelred-stained agarose gels. Right: quantification of percent spliced in (PSI); n = 3. Data information: In d, statistical analysis was performed using t-tests, and the error bars indicate mean ± SD for three independent experiments. ∗∗*p* < 0.01; ∗∗∗*p* < 0.001; ∗∗∗∗*p* < 0.0001.

Of note, the predominant proportion of altered AS events are SE events (Fig. 4a, Table S2). To assess how *THUMPD2* depletion affects SE events, we sought to analyze whether these donor or acceptor splice sites were enriched for special motifs. The results showed canonical sequences of 5’ and 3’ splice sites of major spliceosome-GU and AG-are enriched in the donor and acceptor splice sites, respectively. Notably, the decreased SE donor sites are more enriched for G at +5 while the decreased SE acceptor sites are less enriched for C at -3 (Fig. S11). Suggested THUMPD2 might regulate SE for genes by recognize specific motifs surrounding acceptor/donor sites.

The pre-mRNA targets of changed SE events comprise 956 protein-coding genes (Table S3) contains decreased or increased skipped exons genes (Fig. 4b). These genes are involved in many aspects of life processes, such as *CDK2, CEP131, ENSA, CENPK*, and *CDK20* in cell cycle and proliferation regulation, *ALCAM, MAPK8, LLGL1, FLNA*, and *CD46* in cell signaling pathways, and *NSMF, FKBP15, SLIT2, SETD5*, and *ROBO3* in neuronal development. Notably, a moderate decrease in cell proliferation was observed consistently in both *THUMPD2* KO HEK293T and Hela cells compared with WT cells (Fig. S12, a and b). *THUMPD2* KO induced an increased ratio of G0/G1 phase cells and decreased the ratio of G2/M phase in both cells in HEK293T and Hela cells (Fig. S12, c and d).

We next validate some representative AS events by RT-PCR. For example, the Sashimi plot assays showed that exon skipping in *FLNA, CEP131*, and *FKBP15* and intron retention in *SPOUT1* (Fig. 4c) were much more present in both *THUMPD2* KO compared to WT HEK293T cells, and the increased exon inclusions or intron retention of these mRNAs were confirmed by RT-PCR (Fig. 4d).

Thus, THUMPD2 affects the splicing of a very large number of endogenous pre-mRNAs in many aspects of life processes.

### *THUMPD2* KO upregulates nonsense-mediated mRNA decays

To analyze the effect of THUMPD2 depletion on gene expression, we performed RNA-seq analysis upon *THUMPD2* KO. 511 and 592 genes were up-and down-regulated, respectively (significantly over 2-fold) (Table S4,Fig 5a). GO terms of the upregulated genes revealed several distinct gene clusters (Fig. 5b) enriched for terms including “translation initiation”, “transcriptional activation”, and “nonsense-mediated mRNA decay (NMD) pathway”. The upregulated expression of the transcriptional activation-and translation initiation-related genes in *THUMPD2* knockout cells were confirmed by RT-qPCR assays (Fig. 5c).

**Fig. 5.**
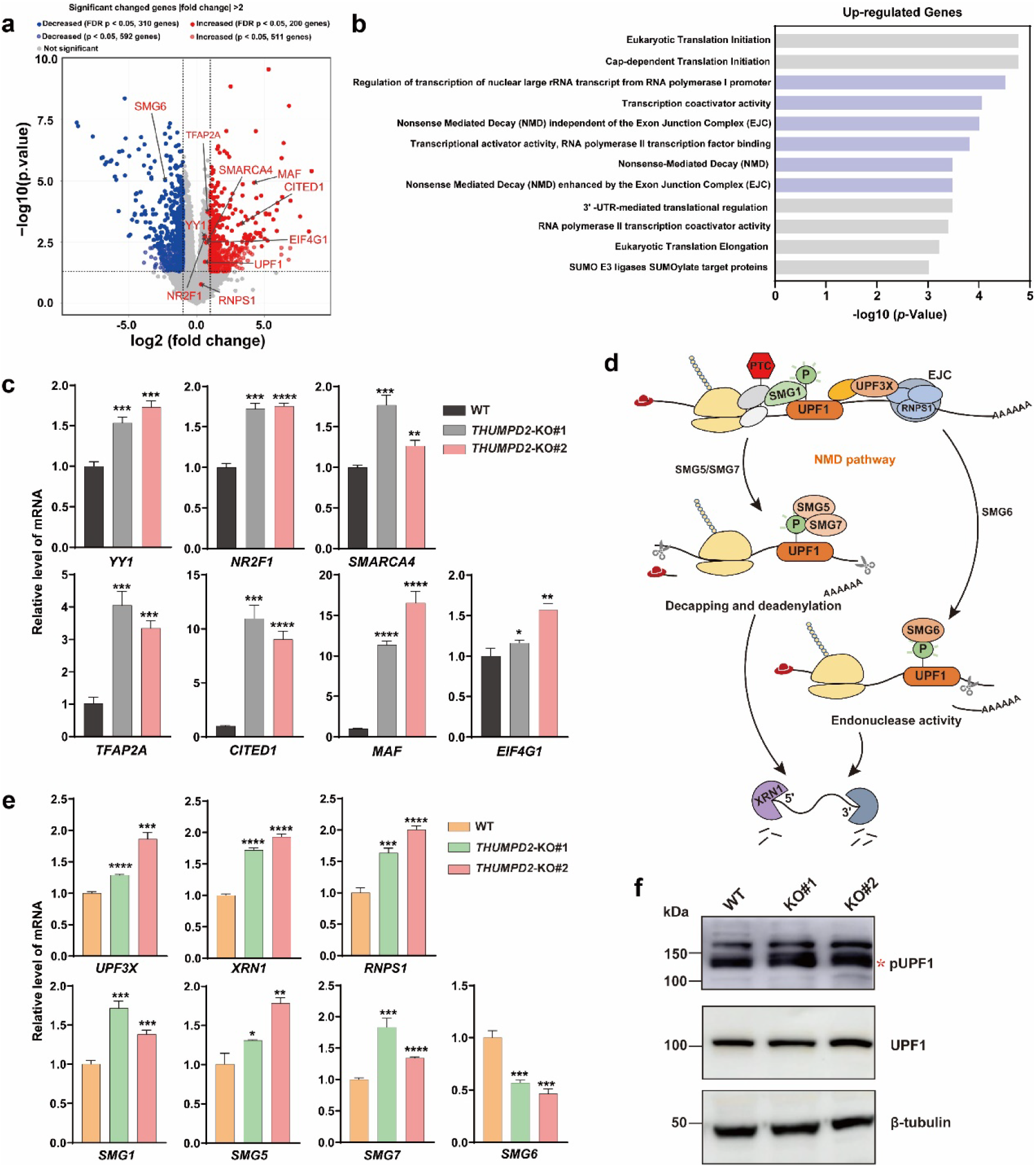
*THUMPD2* KO up regulates the expression level of key factors in NMD pathway. (a) Volcano plot of differential expressed gene analysis results for RNA-seq data comparing *THUMPD2* KO HEK293T cells (n = 4) to control cells (n = 2), the dots for significant regulated genes (|fold change| > 2, *p* < 0.05) were colored in red (increased) and blue (decreased), genes significant but did not passed FDR of 5% were transparent. The dots and labels of interesting genes with transcriptional activation and nonsense-mediated RNA decay (NMD) pathway have been colored in red. (b) Functional categories of upregulated genes in *THUMPD2* KO HEK293T cells showing the *p*-value for the enrichment of biological process GO term. (c) The mRNA level of the key genes involved in the transcription activation-and translation initiation-related genes (*YY1, NR2F1, SMARCA4, TFAP2A, CITED1, MAF*, and *EIF4G1*) were measured by RT-qPCR. Data were normalized to β-actin expression. (d) The schematic diagram of NMD pathway. The key factors involved in NMD pathway were marked. (e) The mRNA level of the key genes involved in the NMD pathway (*UPF3X, XRN1, RNPS1, SMG1, SMG5, SMG7*, and *SMG6*) were measured by RT-qPCR. Data were normalized to β-actin expression. (f) Analysis the protein phosphorylation level of UPF1(pUPF1) by western blotting. Data information: In c and e, statistical analysis was performed using t-tests, and the error bars indicate mean ± SD for three independent experiments. ∗*p* < 0.05;∗∗*p* < 0.01; ∗∗∗*p* < 0.001; ∗∗∗∗*p* < 0.0001.

The NMD pathway is a cellular RNA surveillance mechanism for selectively degrading transcripts with premature termination codons (PTCs) in ∼ 55 nt upstream of exon junction complex (EJC). It has been reported that approximately one third of AS events can produce transcripts containing PTCs that trigger NMD. Recalling the abundant AS events that we detected in *THUMPD2* KO cells (Fig. 4a), we next examined NMD pathway. The central components of NMD pathway are UPF1 with the stimulated factor -SMG1 and EJC (the core of EJC is joined by other proteins including UPF3X and RNPS1), the downstream key effectors include SMG5/SMG7 or SMG6, and 5’-3’ exoribonuclease XRN1 (*28*) (Fig. 5d). The key event in NMD is the phosphorylation and activation of UPF1 by SMG1. Once UPF1 is phosphorylated, it inhibits translation initiation and recruits factors mediating RNA decay (*29*). Therefore, we analyzed the expression levels of genes mentioned above and the phosphorylation level of UPF1. Except SMG6, the mRNA levels of other NMD key factors were all increased obviously in *THUMPD2* KO than WT cells (∼1.3 to 2-fold changes) (Fig. 5e). Consistently, the phosphorylation level of UPF1 is much higher in both *THUMPD2* KO than WT cells (Fig. 5f). These data support that NMD pathway activity is obviously up regulated upon *THUMPD2* KO. Thus, it suggests that the abundant changed AS events caused by loss of THUMPD2 further elicit NMD pathway.

## Discussion

From yeast to human, the increased number of splicing events is coupling with elevated number of modifications on snRNAs and regulatory splicing factors (*11*). Previous studies have shown that snRNAs and m^6^A modifications could enhance the efficiency and fidelity of pre-mRNA splicing (*30, 31*), but the specific mechanisms underlying how RNA modifications modulate pre-mRNA splicing remain unclear. Specially, m^2^G72 is absent in lower eukaryotes but emerged in vertebrates on U6 snRNA of the splicing catalysis center. Our data show that this U6 m^2^G72 increases the pre-mRNA splicing activity of the major spliceosome. The U6 ISL, which constitutes the main part of the splicing catalytic center, is specified by the bulged secondary structure. The conformation of this RNA bulge is known to be essential for the coordination of the two catalytic metal ions M1 and M2, as well as for both steps of the splicing reaction (*7*). Since G72 directly coordinates M1, a subtle change introduced by m^2^G could plausibly impact splicing catalysis. Structurally, G72 base-paring with C62 (flanked by bulged U74) is positioned within a continuous stack of base-paired nucleotides. A recent study of tRNA^Trp^ (*32*) reported that modification of G with a methyl group (added to the N2 atom) enhances hydrophobicity and thereby strengthens base-stacking interaction (*33, 34*). It is also conceivable that the m^2^G72 modification may stabilize the conformation of the splicing center by chelating M1 to facilitate splicing catalysis.

*THUMPD2* KO leads to a total of 4113 significant alternative splicing events that affects around two thousand pre-mRNA species in HEK293T cells, suggesting a wide range of targets of splicing regulation by THUMPD2. These altered-splicing of numerous genes are involved in a variety of general biological processes, including cell cycle, cell proliferation, cell signaling pathways, neuronal development, and apoptosis. Consistently, *THUMPD2* depletion further caused moderately decreased cell proliferation in human cells. One-third of pre-mRNA AS events in human and mouse cells routinely introduce PTCs to trigger NMD (*35, 36*). Upon *THUMPD2* depletion, abundant AS events were observed, thus NMD pathway was consequently upregulated. Our findings suggest that THUMPD2 acts as a robust regulator in global pre-mRNA splicing.

## Methods

### Cell culture

HEK293T, SH-SY5Y, Hep G2, HeLa, U-2 OS, SK-BR-3, MDA-MB-231, MCF-7, NCl-H1299, A549, HCC38, RPE, and BT-549 cells were purchased from the cell resource center of the Shanghai Institutes for Biological Sciences, Chinese Academy of Sciences, Shanghai, China. Except for RPE and SH-SY5Y, all of them were cultured at 37°C with 5% CO_2_ in Dulbecco’s modified Eagle’s medium (DMEM, Corning) supplemented with 10% fetal bovine serum (Gibco) and 1% penicillin-streptomycin. The medium for RPE is Dulbecco’s Modified Eagle Medium/Nutrient Mixture F-12 (DMEM/F-12, Gibco). The medium for SH-SY5Y is equal mixture of DMEM/F-12 and DMEM. The viable cell numbers were counted by 0.4% trypan blue staining assays. Insect cells, *Spodoptera frugiperda* Sf9 and High Five, were cultured on a shaking incubator at 27°C and 120 rpm in ESF921 medium (Expression Systems).

### Construction of knockout cell lines

Sense and anti-sense oligonucleotides for a guide RNA (sgRNA) were computationally designed for the selected genomic targets (http://crispor.tefor.net) and were cloned into pX330-mcherry vector (Addgene, 98750) which expresses red fluorescence protein. Three sgRNA sets were designed for *THUMPD2*. The sgRNA sequences (5’ to 3’) for relevant genes and targeting sites in HEK293T, HeLa, and HeLa S3 cells are shown in Fig. S4, a, b, and c, respectively. For generating KO cell lines, sgRNA plasmids were transfected into cells using EZ Cell transfection reagent. After transfection for 36 h, cells expressing red fluorescent protein were enriched by FACS Aria Fusion SORP (BD Bioscience) and plated into a dish at a very low density. After around 14 days, single colonies were picked and plated into a well of a 96-well plate. Genotype of the stable cell lines was identified by confirming the frameshift mutations in the target region of genome.

### Western blotting

Cell lysates and immunoprecipitation complexes were separated by sodium dodecyl sulfate-polyacrylamide gel electrophoresis (SDS-PAGE), and the protein bands were transferred to 0.2 µm or 0.45 µm PVDF membranes. After blocking with 5% (w/v) non-fat dried milk, the membranes with targeted proteins were incubated with the corresponding primary antibodies overnight at 4°C. Membranes were then washed three times with PBST (phosphate-buffered saline with Tween-20) and incubated with HRP-conjugated secondary antibody at room temperature for 1 h. After washing three times with PBST, the membranes were treated with the chemiluminescent substrates (EpiZyme), and imaging was performed using the Amersham™ ImageQuant™ 800 (GE Healthcare).

### Confocal immunofluorescence microscopy

HEK293T cells were transfected with pcDNA3.1(+)-THUMPD2-HA plasmid. After transfection for 36 h, the cells were fixed in 4% paraformaldehyde for 10 min and then permeated in 0.2% Triton X-100 for 5 min on ice. After washing with phosphate-buffered saline (PBS), the fixed cells were blocked in PBS containing 5% bovine serum albumin (BSA) and then incubated with mouse anti-HA antibody (6E2, Cell signaling Technology) with 1:800 dilution overnight at 4°C. The cells were then immunolabeled with Alexa Fluor 647-conjugated donkey anti-mouse IgG (Yeasen, Shanghai) in PBS with 1:400 dilution for 2 h and the nuclear counterstain DAPI (Thermo Fisher Scientific) for 5 min at room temperature. Fluorescent images were taken and analyzed using a LSM980 Airyscan2 confocal microscope (Zeiss).

### Immunoprecipitation

The cDNA of *HsTHUMPD2* (NM_025264.5) (along with a DNA sequence encoding a C-terminal HA-tag) and the cDNA of *HsTRMT112* (NM_016404.3) (along with a DNA sequence encoding a C-terminal Flag-tag) was inserted in pcDNA3.1(+), respectively. For immunoprecipitation, HEK293T cells were transfected with pcDNA3.1(+)-THUMPD2-HA or pcDNA3.1(+)-TRMT112-Flag. EZ Cell transfection reagent was used for transfection according to the manufacturer’s protocol. After transfection for 36 h, the cells were washed three times with ice-cold PBS and then lysed with ice-cold lysis buffer (50 mM Tris-HCl (pH 7.4), 150 mM NaCl, 1 mM EDTA, 1% NP-40, and 0.25% deoxycholic acid) supplemented with a Proteinase Inhibitor Cocktail (MedChemExpress) for 20 min. The supernatant was collected by centrifugation at 12,000 rpm for 10 min. Subsequently, the supernatant was incubated with the anti-HA magnetic beads or anti-Flag magnetic beads (MedChemExpress) with gentle agitation for appropriate time. Recovered immunoprecipitation complexes were washed three times with ice-cold lysis buffer. All procedures were performed at 4°C. The immunoprecipitation complex was then eluted by buffer (150 mM NaCl, 20 mM Tris-HCl, pH 7.5, and 1% SDS). The eluted IP samples were loaded on SDS-PAGE followed by Western blot analysis. To identify the protein interactome of *Hs*THUMPD2, the eluted samples of HA-THUMPD2 were digested into peptides and then subjected to LC-MS/MS for protein identification.

### Preparation of snRNA transcripts

The DNA sequences of the T7 promoter and the U6 snRNAs (of *Homo sapiens, Mus musculus*, and *Danio rerio*) or *Hs*U6atac snRNA were obtained from NCBI and cloned into pTrc99b vector (two restriction enzyme cutting sites for the vector were EcoRI and BamHI) to construct pTrc99b-T7-snRNA plasmids. The sequences of mutated *Hs*U6snRNA including U6-loop(atac), U6-d5’SL, U6-d3’U-tail, U6-telestem, U6 ISL-apical loop-m.A(G65U), U6 ISL-apical loop-m.B(C66G), U6 ISL-apical loop-m.C (G67A), U6 ISL-apical loop-m.D(C68A), U6 ISL-apical loop-m.E(A69C), U6 ISL-apical loop-m.F(C61U), U6 ISL-apical loop-m.G(A73G), and U6 ISL-apical loop-m.H(dU) were constructed into pTrc99b vector as well as wildtype U6 snRNA. All snRNA transcripts were generated via *in vitro* transcription using T7 RNA polymerase. The transcribed snRNAs were denatured and annealed to form the right conformation in 5 mM magnesium chloride (MgCl_2_).

### Cell Counting Kit-8 (CCK-8) assays

Cell proliferation was determined using the Cell Counting Kit-8. Briefly, 1.2 x 10^3^ WT or *THUMPD2* knockout HEK293T or HeLa cells were seeded in a 96-well flat-bottomed plate under normal culture. At days 1, 2, 3, 4, and 5, 10 µl of CCK-8 reagent (Vazyme) was added into each well and the cells were incubated for 2 h at 37°C with 5% CO_2_. The optical density at 450 nm (OD450) was measured using a Spark multimode plate reader (Tecan) to assay the growth curve. The experiments were repeated for three times.

### snRNA methyl-transfer assays

To confirm the modification introduced on snRNAs by purified *Hs*THUMPD2, *Hs*TRMT112, *Hs*THUMPD2-TRMT112, *Mm*THUMPD2-TRMT112, and *Dr*THUMPD2-TRMT112 is indeed m^2^G, the reactions were carried out at 37°C for 2 h in a 50 µl reaction mixture containing 50 mM Tris-HCl (pH 7.5), 100 mM NaCl, 5 mM MgCl_2_, 2 mM DTT, 200 µM SAM, 5 µM snRNA, and 1 µM THUMPD2-TRMT112 complex, respectively. After the reaction, the snRNAs were extracted with phenol/chloroform and precipitated using a two-fold volume of ethanol. Subsequently, the tRNAs were digested and then subjected to UPLC-MS/MS analysis to detect and quantify m^2^G as described above.

### Construction of lentivirus mediated *THUMPD2* expression cell lines

Lentiviruses were generated in HEK293T cells by EZ Cell transfection reagent co-transfection of lentiviral-based pLVX-IRES-PURO plasmids, psPAX2(lentiviral packaging plasmid), and pMD2.G(VSV-G envelope expressing plasmid) for 48h. THUMPD2 knockout and WT HEK293T or HeLa S3 cells were incubated with viruses of *HsTHUMPD2*-HA and *HsTHUMPD2*(D329A)-HA, for 24 h respectively. Followed by culturing in fresh media for 24 h, cells were then cultured in media supplemented with 2 µg/ml puromycin (MedChemExpress) for another 96 h.

### Statistical analysis

Student’s t test was performed to compare the differences between knockout groups relative to their controls using GraphPad Prism v8. All computational results were presented as the mean ± SD. *p*-values (*p*) are indicated in the figures above the two groups.

## Acknowledgments

This work is supported by National Key Research and Development Program of China [2021YFA1100800, 2020YFA0803400]; National Natural Science Foundation of China [32022040, 31971230]; This work is also supported by the Shanghai Frontiers Science Center for Biomacromolecules and Precision Medicine in ShanghaiTech University.

We thank the Molecular and Cell Biology Core Facility (MCBCF), the Multi-Omics Core Facility (MOCF), and the Molecular Imaging Core Facility (MICF) at the School of Life Science and Technology, ShanghaiTech University for providing technical support. We also thank the Analytical Chemistry platform (ShanghaiTech University, SIAIS) for technical assistance with protein identification by MS. We thank Prof. Hong Cheng and Jing Yi Hui at Shanghai Institute of Biochemistry and Cell Biology, Chinese Academy of Sciences for technical assistance.

## References

1. M. S. Jurica, M. J. Moore, Pre-mRNA splicing: awash in a sea of proteins. Molecular cell 12, 5–14 (2003).

2. M. E. Wilkinson, C. Charenton, K. Nagai, RNA Splicing by the Spliceosome. Annual review of biochemistry 89, 359–388 (2020).

3. R. Wan, R. Bai, X. Zhan, Y. Shi, How Is Precursor Messenger RNA Spliced by the Spliceosome? Annual review of biochemistry 89, 333–358 (2020).

4. S. Valadkhan, The spliceosome: a ribozyme at heart? Biological chemistry 388, 693–697 (2007).

5. A. L. Didychuk, S. E. Butcher, D. A. Brow, The life of U6 small nuclear RNA, from cradle to grave. RNA 24, 437–460 (2018).

6. C. J. McManus, M. L. Schwartz, S. E. Butcher, D. A. Brow, A dynamic bulge in the U6 RNA internal stem-loop functions in spliceosome assembly and activation. RNA 13, 2252–2265 (2007).

7. Y. Lee, D. C. Rio, Mechanisms and Regulation of Alternative Pre-mRNA Splicing. Annual review of biochemistry 84, 291–323 (2015).

8. M. Frye, B. T. Harada, M. Behm, C. He, RNA modifications modulate gene expression during development. Science 361, 1346–1349 (2018).

9. R. Reddy, D. Henning, G. Das, M. Harless, D. Wright, The capped U6 small nuclear RNA is transcribed by RNA polymerase III. The Journal of biological chemistry 262, 75–81 (1987).

10. A. Moenne et al., The U6 gene of Saccharomyces cerevisiae is transcribed by RNA polymerase C (III) in vivo and in vitro. The EMBO journal 9, 271–277 (1990).

11. P. Morais, H. Adachi, Y. T. Yu, Spliceosomal snRNA Epitranscriptomics. Frontiers in genetics 12, 652129 (2021).

12. P. Epstein, R. Reddy, D. Henning, H. Busch, The nucleotide sequence of nuclear U6 (4.7 S) RNA. The Journal of biological chemistry 255, 8901–8906 (1980).

13. F. Harada, N. Kato, S. Nishimura, The nucleotide sequence of nuclear 4.8S RNA of mouse cells. Biochemical and biophysical research communications 95, 1332–1340 (1980).

14. K. E. Pendleton et al., The U6 snRNA m^6^A Methyltransferase METTL16 Regulates SAM Synthetase Intron Retention. Cell 169, 824–835.e814 (2017).

15. M. Mendel et al., Splice site m^6^A methylation prevents binding of U2AF35 to inhibit RNA splicing. Cell 184, 3125–3142.e3125 (2021).

16. S. M. Fica et al., RNA catalyses nuclear pre-mRNA splicing. Nature 503, 229–234 (2013).

17. T. A. Steitz, J. A. Steitz, A general two-metal-ion mechanism for catalytic RNA. Proceedings of the National Academy of Sciences of the United States of America 90, 6498–6502 (1993).

18. A. A. Patel, J. A. Steitz, Splicing double: insights from the second spliceosome. Nature reviews. Molecular cell biology 4, 960–970 (2003).

19. S. K. Purushothaman, J. M. Bujnicki, H. Grosjean, B. Lapeyre, Trm11p and Trm112p are both required for the formation of 2-methylguanosine at position 10 in yeast tRNA. Molecular and cellular biology 25, 4359–4370 (2005).

20. X. Wang et al., Structural basis of N(6)-adenosine methylation by the METTL3-METTL14 complex. Nature 534, 575–578 (2016).

21. S. Kadaba et al., Nuclear surveillance and degradation of hypomodified initiator tRNAMet in S. cerevisiae. Genes & development 18, 1227–1240 (2004).

22. G. Bourgeois, J. Létoquart, N. van Tran, M. Graille, Trm112, a Protein Activator of Methyltransferases Modifying Actors of the Eukaryotic Translational Apparatus. Biomolecules 7, 7 (2017).

23. W. Q. Yang et al., THUMPD3-TRMT112 is a m^2^G methyltransferase working on a broad range of tRNA substrates. Nucleic acids research 49, 11900–11919 (2021).

24. S. Boonanuntanasarn, S. Panyim, G. Yoshizaki, Characterization and organization of the U6 snRNA gene in zebrafish and usage of their promoters to express short hairpin RNA. Marine genomics 1, 115–121 (2008).

25. J. S. Sun, J. L. Manley, The human U6 snRNA intramolecular helix: structural constraints and lack of sequence specificity. RNA 3, 514–526 (1997).

26. X. Zhan, C. Yan, X. Zhang, J. Lei, Y. Shi, Structure of a human catalytic step I spliceosome. Science 359, 537–545 (2018).

27. S. Shen et al., rMATS: robust and flexible detection of differential alternative splicing from replicate RNA-Seq data. Proceedings of the National Academy of Sciences of the United States of America 111, E5593–5601 (2014).

28. S. L. Wolin, L. E. Maquat, Cellular RNA surveillance in health and disease. Science 366, 822–827 (2019).

29. M. W. Popp, L. E. Maquat, Organizing principles of mammalian nonsense-mediated mRNA decay. Annual review of genetics 47, 139–165 (2013).

30. Y. Ishigami, T. Ohira, Y. Isokawa, Y. Suzuki, T. Suzuki, A single m^6^A modification in U6 snRNA diversifies exon sequence at the 5’ splice site. Nature communications 12, 3244 (2021).

31. G. Dönmez, K. Hartmuth, R. Lührmann, Modified nucleotides at the 5’ end of human U2 snRNA are required for spliceosomal E-complex formation. RNA 10, 1925–1933 (2004).

32. A. Hirata et al., Distinct Modified Nucleosides in tRNA^Trp^ from the Hyperthermophilic Archaeon Thermococcus kodakarensis and Requirement of tRNA m^2G10/m22G10^ Methyltransferase (Archaeal Trm11) for Survival at High Temperatures. Journal of bacteriology 201, e00448–19 (2019).

33. T. Hermann, D. J. Patel, RNA bulges as architectural and recognition motifs. Structure (London, England : 1993) 8, R47–54 (2000).

34. S. L. Ginell, R. Parthasarathy, Conformation of N2-methylguanosine, a modified nucleoside of tRNA. Biochemical and biophysical research communications 84, 886–894 (1978).

35. T. Kurosaki, M. W. Popp, L. E. Maquat, Quality and quantity control of gene expression by nonsense-mediated mRNA decay. Nature reviews. Molecular cell biology 20, 406–420 (2019).

36. B. P. Lewis, R. E. Green, S. E. Brenner, Evidence for the widespread coupling of alternative splicing and nonsense-mediated mRNA decay in humans. Proceedings of the National Academy of Sciences of the United States of America 100, 189–192 (2003).

